# Brain map of aging-induced alterations in membrane ganglioside pattern

**DOI:** 10.1101/2023.08.30.555520

**Authors:** Durga Jha, Gabriela Dovrtelova, Petr Telensky, Seweryn Olkowicz, Katerina Coufalikova, Lukas Opalka, Karel Kubicek, Bára Černá, Roman Hajek, Silvie Belaskova, Jana Klanova, Ales Hampl, Jiri Damborsky, Zdenek Spacil

## Abstract

Recent efforts to develop comprehensive metabolome and proteome brain atlases for animal models have yielded significant progress. However, the ganglioside (GSs) profile of these models remains largely unexplored. As essential components of the brain, GSs play a crucial role in neuronal function. To address this gap in knowledge, we conducted an in-depth analysis of the young and adult rat brain, as well as its major brain regions, using ultra-high performance liquid chromatography and tandem mass spectrometry in positive and negative ion modes. We also analyzed GSs in cerebrospinal fluid (CSF) and serum from matched samples. Our findings indicate a shift in the ratio of a-series to b-series GSs with age, along with region-specific changes accompanied by aging. This could improve our understanding of brain aging and neurodegenerative diseases. Our study complements existing brain atlases of lipidome and protein expression and highlights the importance of further investigating the mechanisms underlying these GSs changes and the potential therapeutic implications of our findings.

## Introduction

Gangliosides (GSs) are sialylated glycosphingolipids enriched explicitly in the CNS, composing up to ten percent of the brain’s lipidome^1^. They consist of a hydrophobic ceramide chain embedded in the plasma membrane and an oligosaccharide moiety with sialic acid extended in the extracellular space. The carboxyl group of sialic acid exists predominantly in ionic form (hydrophilic) in lipid bilayers^2^. Together with a sphingosine long-chain base component (hydrophobic), they modulate the membrane’s organization by affecting its fluid properties, charge density, surface electrical potential, and pH^3^. Besides their role in maintaining neuronal membrane integrity, GSs are also involved in cell signaling and cell differentiation^3^. Their synthesis usually initiates in the luminal membrane of the endoplasmic reticulum and begins with the addition of the sialic acid to a lactosylceramide to form GM3. GM3 and GD3 are the precursors for the a-and-b-series GSs, respectively. GM3 and GD3 are referred to as the simple GSs. The primary complex mammal brain GSs are GM1, GD1a, GD1b, and GT1b (Figure 1)^4^. These complex GSs occur through a collection of enzymes such as sialyltransferases, galactosyl transferases, and N-galactosaminyltransferases^5^. They are localized in specific membrane microdomains, and their composition could regulate the proteins responsible for signal transduction and cell metabolism ^6^. Thus, their profiles can signal the metabolic changes in neuronal cells, such as aging and neurodegenerative disorders.

**Figure 1.**
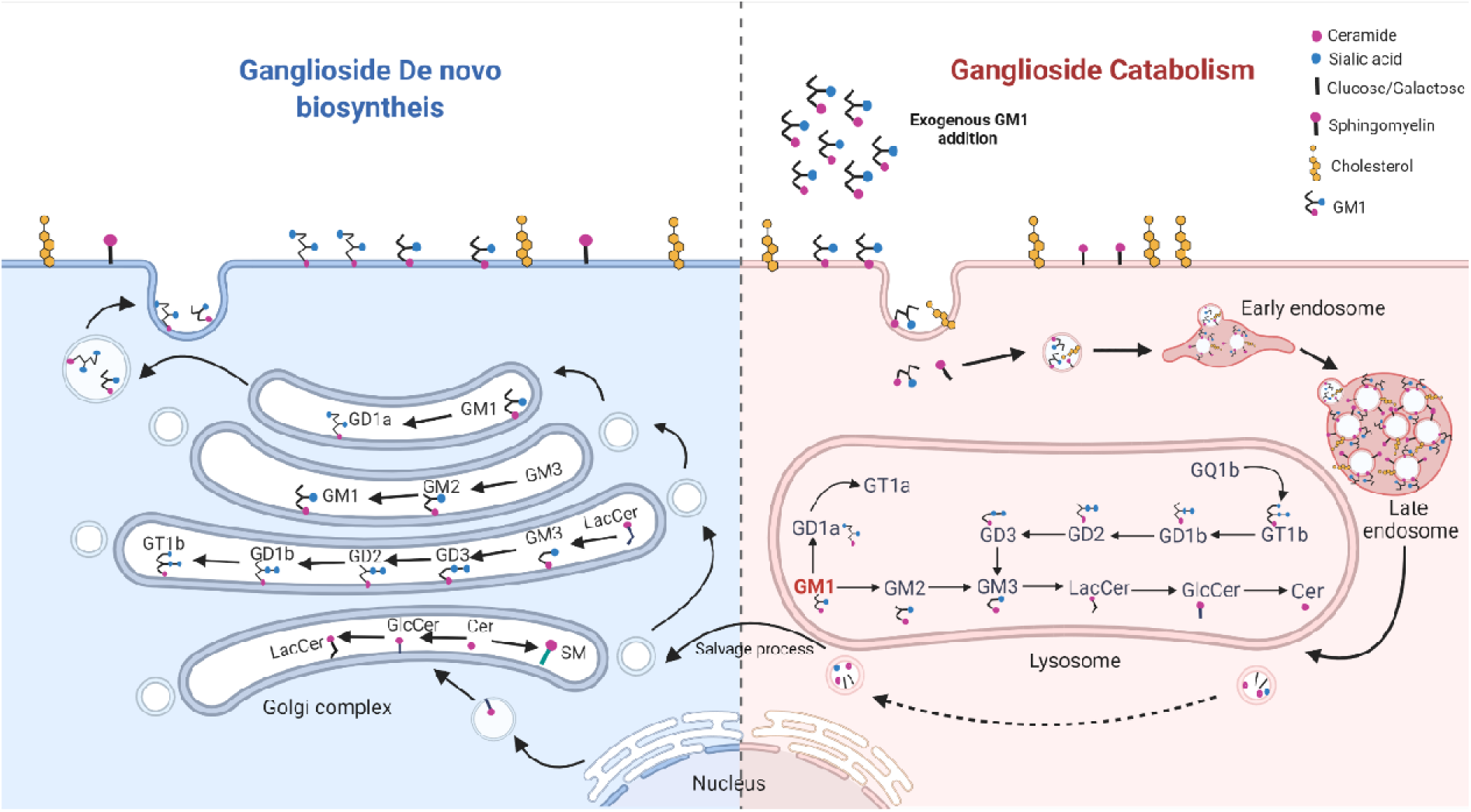
Intracellular metabolism of GSs. GSs catabolism begins with endocytosis in endosomes and their movement to the lysosomes, where they are broken into their constituent parts, which can be used in the salvage process for the de novo biosynthesis. The synthesis begins with ceramide synthesis in the endoplasmic reticulum. The lactosylceramide is moved to the Golgi complex, where the GSs synthesis occurs and is further transported to the plasma membrane.

Aging is a natural biological process that leads to physiological decline and is a big risk factor for several neurodegenerative diseases such as Alzheimer’s. The altered GSs biosynthesis and metabolism in brain tissue has been reportedly linked with neurodevelopmental defects^7,8^, and changes in GS levels have been implicated as a potential mechanism of the development of neurodegeneration^3,9^.

Owing to the relative complexity of the CNS and marked changes in it during aging, much emphasis has been put on resolving the global proteome and metabolome and understanding how their levels are affected across functionally and morphologically amongst different structures and how the changes in aging modulate their levels^10–13^. Several groups are working towards preparing CNS atlas of rats’ brains using the genomic and metabolomic profiles. However, results on aging-induced alterations in GS profile^14,15^ shows ambiguity compared to the more recent studies ^9,16,17^. There is also insufficient information regarding the quantitative spatial distribution of individual GSs in mammal brain tissue.

The brain is a complex organ with a diverse composition of cellular populations in different structures based on their functions. Thus, it is crucial to study diverse structures to get region-specific GSs profiles. It is also imperative to examine whether the brain changes are reflected concurrently in the associated sample matrices such as cerebrospinal fluid (CSF) and the serum. Here we report a comprehensive map of aging-induced alterations in GS profiles in specific mammalian brain regions, CSF, and serum. We used advanced analytics by ultra-high performance liquid chromatography (UHPLC) and selected reaction monitoring (SRM) tandem mass spectrometry (MS) to perform highly selective and quantitative determination of abundant brain GS species.

## Result

### 1. GSs profiling in the rat brain, CSF, and serum

In order to visualize the complete GS profile in the rat, three different relevant biological matrices were utilized, including the brain, CSF, and serum from the corresponding rats. Since there is a region-based distinct lipidome and proteome expression in the brain, the rat brain tissues were obtained from eleven different anatomical structures to have a global overview of GS presence (Figure 2A). These regions were neocortex (NEO), allocortex (ALLO), cerebellum (CB), striatum STR), olfactory bulb (OB), medial basal forebrain (MF), hypothalamus (HYPO), brainstem (BS), cingulate cortex (CING), ventral (VHIP) and dorsal hippocampus (DHIP).

**Figure 2.**
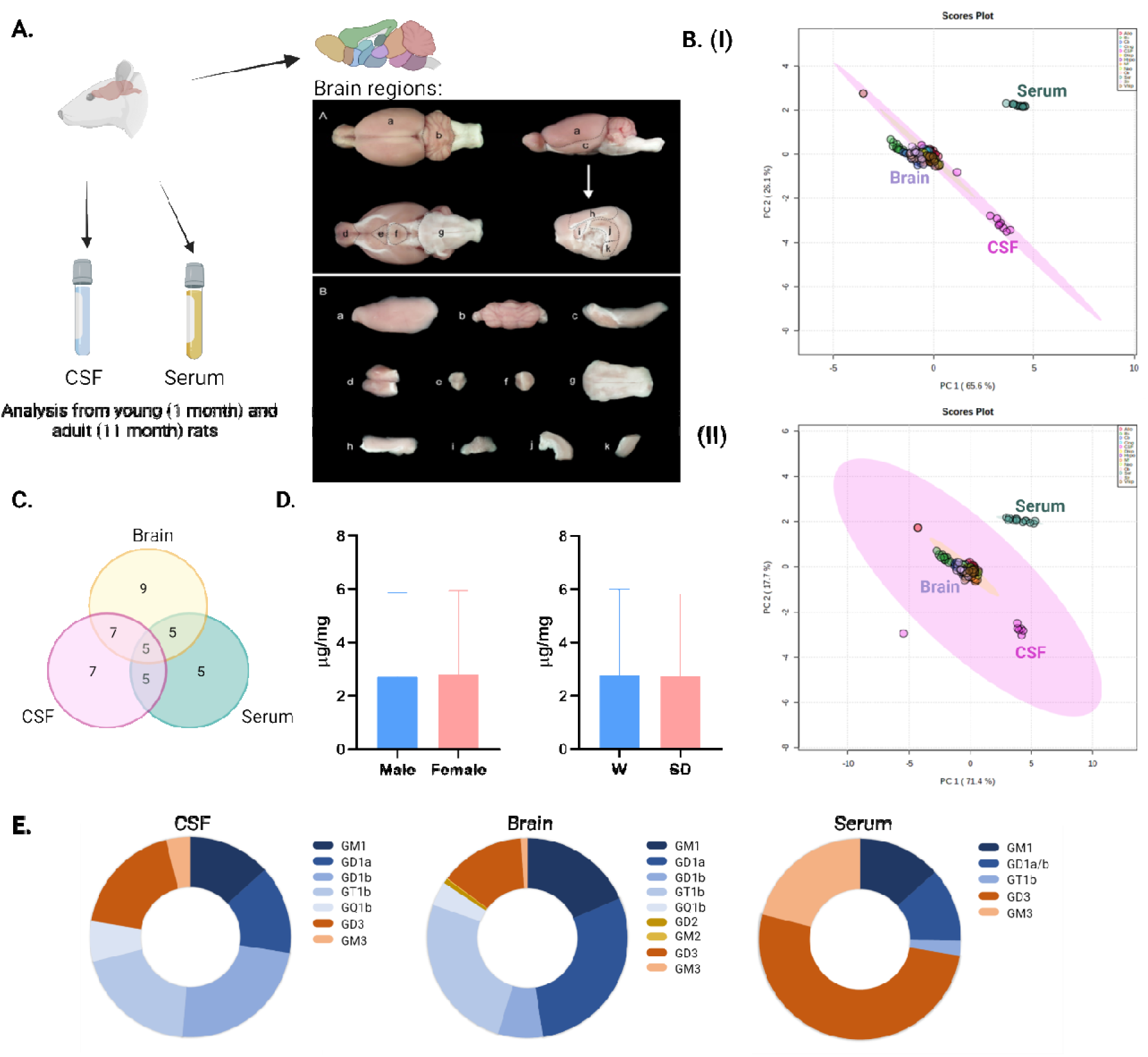
**A.** Analysis was performed in the CSF, serum, and brain tissue collected from young and adult rats. The brain tissue was further dissected in different regions: **Panel A:** The whole brain was partitioned by separating neocortex (a), cerebellum (b), allocortex (c), and olfactory bulbs (d), followed by the dissection of the medial basal forebrain (e), hypothalamus (f) and the brainstem (g). Subsequently, the right hemisphere was separated and dissected into the striatum (i), dorsal (j) and ventral (k) hippocampus, and the cingulate cortex (h). The approximate lines of cut are marked with dashed lines. **Panel B:** Individual dissected brain regions. The labeling of regions is the same as in Panel A. **B. (I)** PCA performed with the complete GS profile observed in young rats’ serum, CSF, and different brain regions segregated based on component 1 and component 2, accounting for 65.6% and 26.1%, respectively. CSF profile completely segregated from the brain and serum. **(II)** PCA performed with the complete GS profile observed in older rats’ serum, CSF, and different brain regions segregated based on components 1 and 2, accounting for 71.4% and 17.7%, respectively. DHIP from the other brain regions. **C.** Venn diagram of the number of GS sub-classes analyzed in different sample types. **D.** GS levels in male and female rats and Wistar and Sprague Dawley rats.

Since the glycosphingolipids are more prevalent in the brain, we profiled nine major sub-classes of GS - GM1, GD1a, GD1b, GT1b, GM3, GD3, GM2, and GD2. As the proximity of the matrix from the brain reduced, the GS classes observed were also reduced with the least number of sub-classes observed in the serum sample (Figure 2B).

A principal component analysis (PCA) was performed to discover distinct patterns in the data. PCA performed with the three sample matrices showed clustering of samples from all brains regions clustered together, whereas the serum samples were markedly separate, suggesting the difference in their GS composition. These changes were reflected in both young and adult rat samples (Figure 2C). The PCA plot also showed biological replicates clustering together with points to excellent reproducibility. Where the brain and CSF were contained over 90% of the complex GSs found abundantly in the neuronal populations - GM1, GD1a, GD1b, GT1b, and GQ1b, the serum had more pronounced levels of GSs more visible in the endothelial cells such as GM3 and GD3, with significantly less presence of complex GSs (Figure 2D).

Before further analysis was performed, it was necessary to ensure that the changes reported here were not affected due to gender or the strains. The overall level of GSs showed no difference between male and female rats and the W and SD rats (Figure 2E). The detailed analysis between the genders and the strains for individual brain regions, CSF and serum, did not produce any differences (Supplementary Figure 1).

### 2. Mapping brain-region specific presence of GSs

The CNS has diverse anatomical regions, with each region carrying different cellular populations and functions. Hierarchical clustering and the PCA (Figure 3a) were performed with the GS quantitative information in the 11 brain regions analyzed. Based on PCA, it is evident that the brain stem, cerebellum, and olfactory bulb were the most distinct brain region in both young and adult rats. The rest of the regions were clustered tightly together.

**Figure 3.**
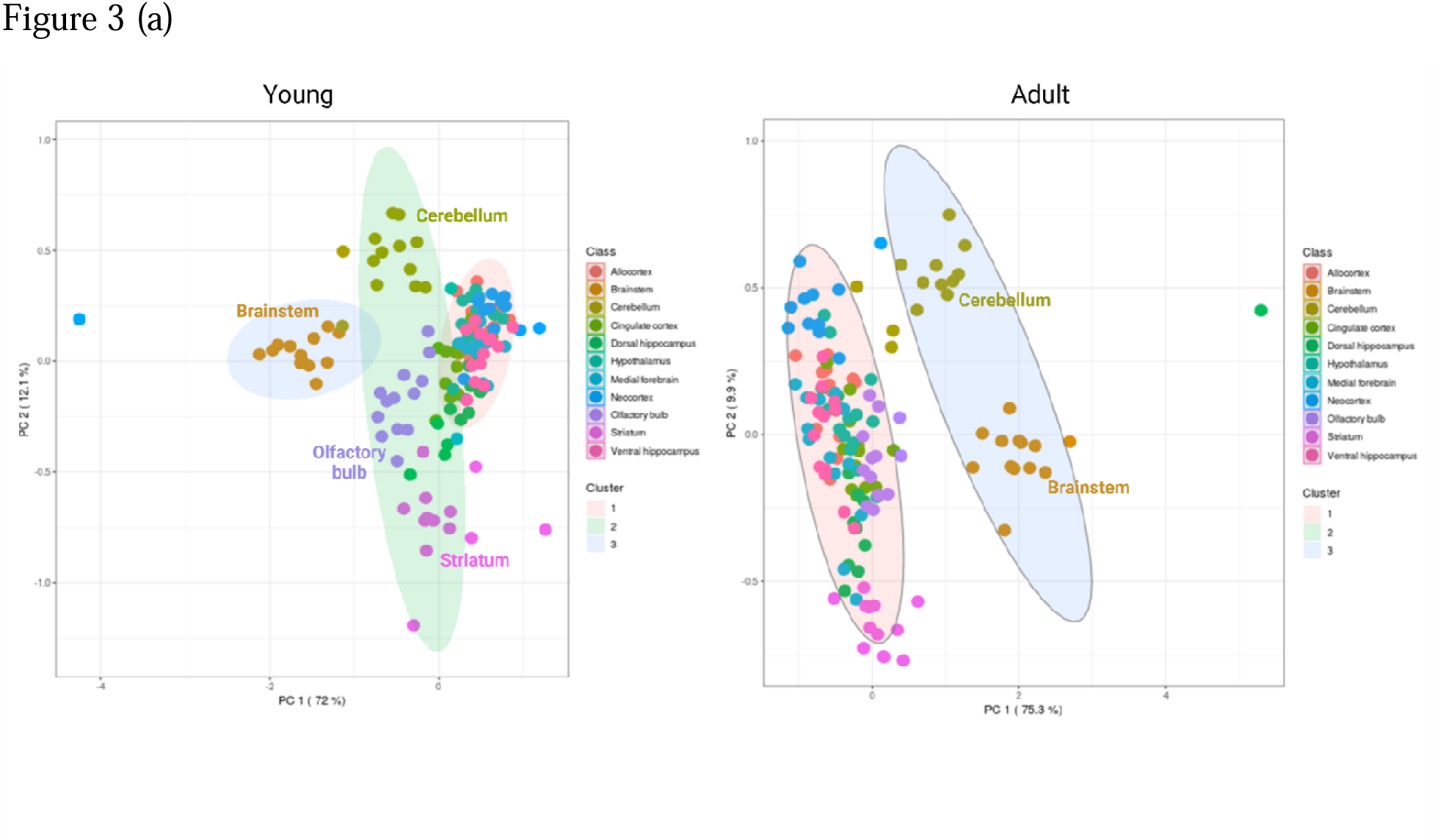
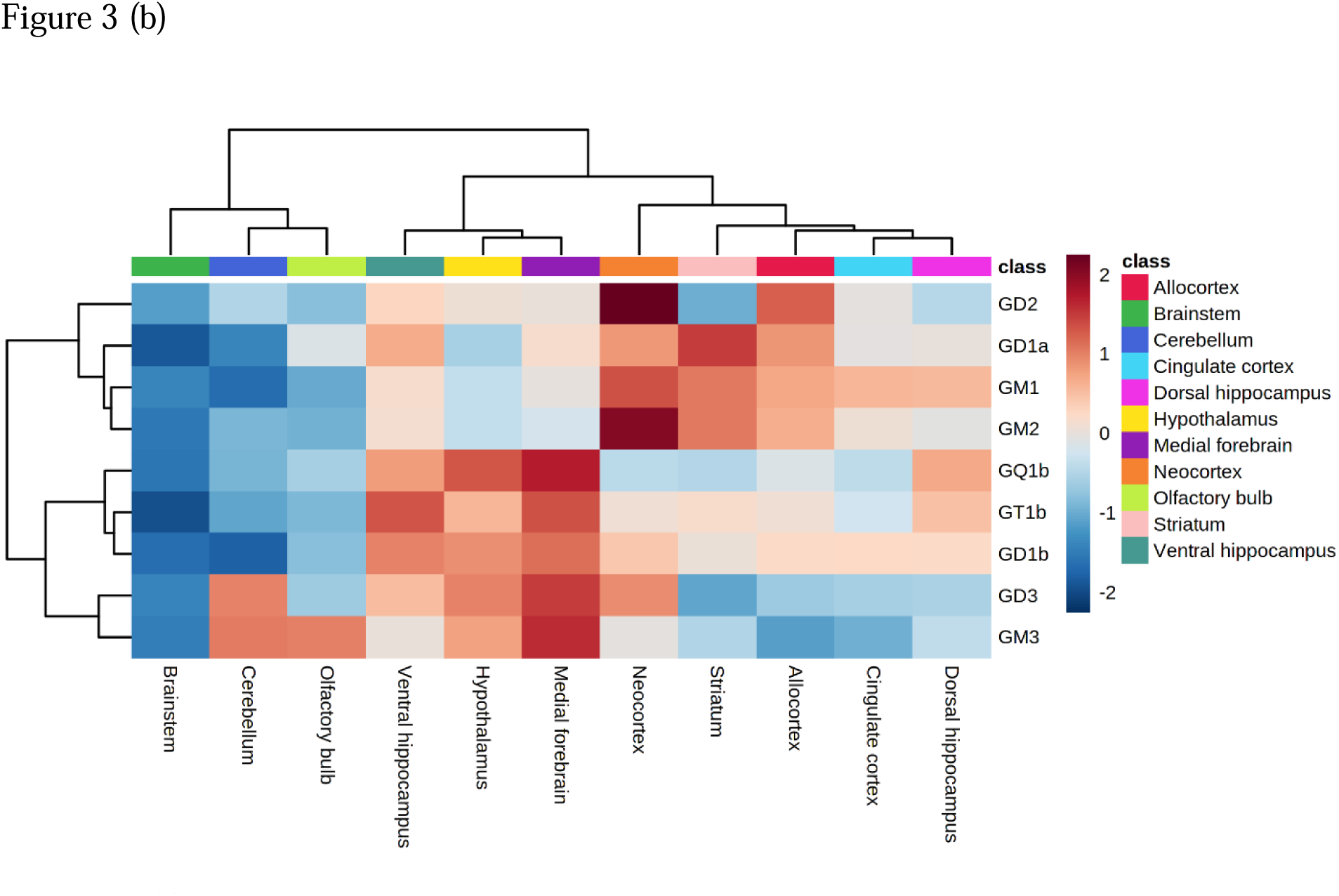
(a) K means a plot of the quantitative data of GS from the eleven brain regions. Three different clusters were formed with brainstem and cerebellum segregating distinctly in older rats, whereas in the younger rats, brainstem, cerebellum, olfactory bulb, and striatum were separated from the rest of the brain regions. (b) Heatmap with hierarchical clustering. The individual biological replicates were averaged to visualize the key groups. A- and B-series GS were clustered in two groups. The brain regions are segregated based on their function and affinity towards a- or b-series GSs.

The hierarchical clustering based on both young and adult rats showed three distinct clusters (Figure 3b). The first cluster was based on the brainstem, cerebellum, and olfactory bulb, also evident in the PCA. These three structures were the least enriched in GSs. Most brain structures comprised 74-90% of complex GSs, whereas the cerebellum and brainstem consisted only 58% of complex GSs, with high levels of simple GSs such as GD3 and GM3. The second cluster wa made of the ventral hippocampus, medial forebrain, and hypothalamus.

Interestingly, these structures were abundant in the b-series GSs such as GD1b, GT1b, GQ1b, and GD3. Finally, the last cluster consisted of the neocortex, cingulate cortex, dorsal hippocampus, allocortex, striatum. Contrary to the second cluster, this group consisted of mostly a-series GSs, especially GM1.

Another intriguing find of this analysis was the separate clustering of the ventral hippocampus and the dorsal hippocampus.

Out of all the complex GSs, the most abundant in most brain structures were GD1a and GT1b, followed by GM1, GT1b and GQ1b, respectively. The least abundant GSs in the brain was GM2.

### 3. Aging relevant changes in the GSs profile

In order to see changes in GSs due to aging, we compared its level in young and adult rats, in all the brain structures, along with CSF and serum. There was a significant decline in polysialic GSs - GD1a, GD1b, GT1b, GQ1b, GD2, and GD3, in almost all structures. On the contrary, there was an increase in the monosialic GSs - GM1, GM2, and GM3 in different regions (Figure 4).

**Figure 4.**
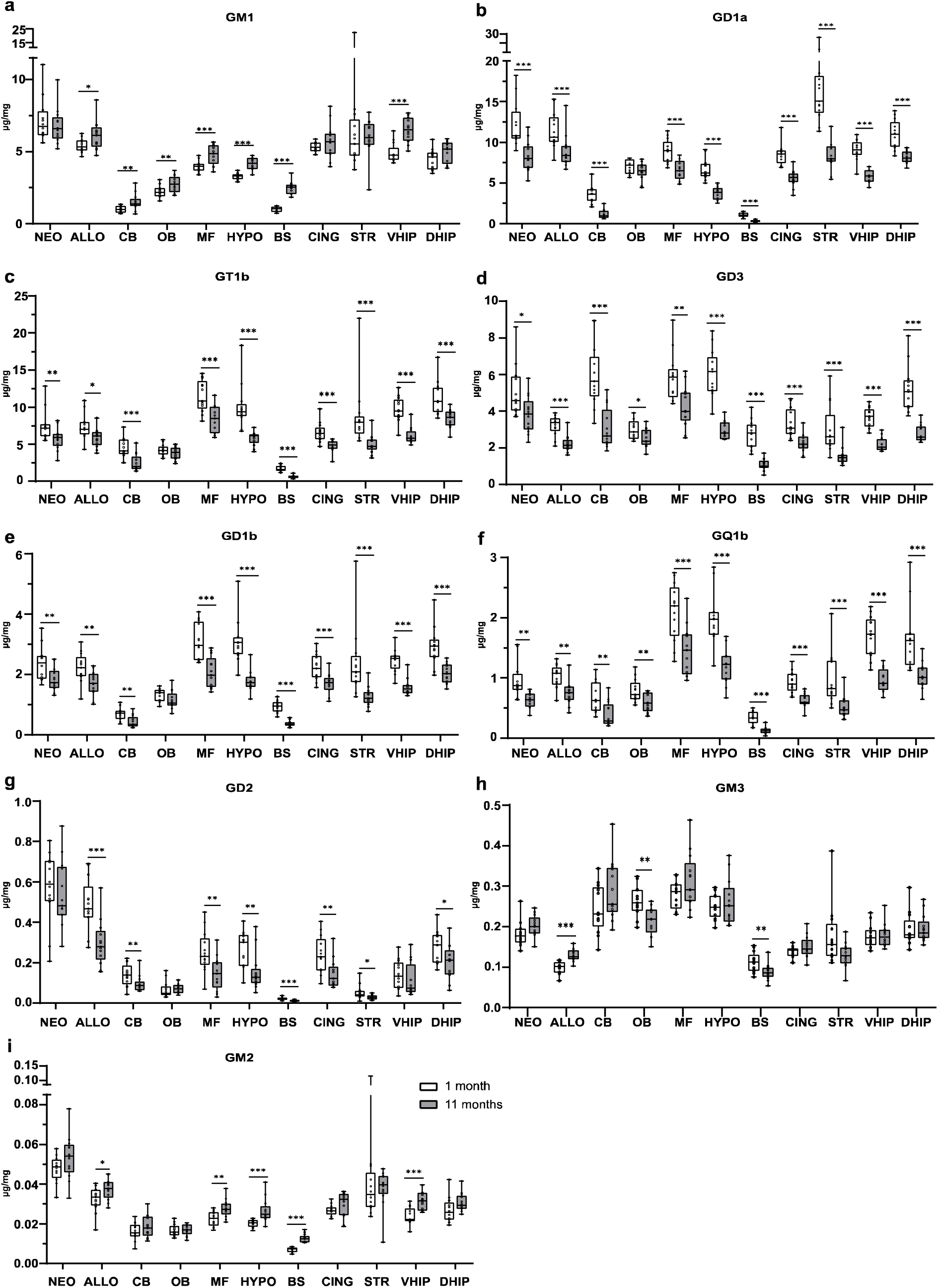
Effect of aging on the rat brain GSs levels in individual rat brain structures. **a–i** Changes in GSs levels (**a** GM1, **b** GD1a, **c** GT1b, **d** GD3, **e** GD1b, **f** GQ1b, **g** GD2, **h** GM3, **i** GM2); ordered by abundance) expressed as µg of GSs per mg of brain tissue DW. The levels were determined in 1 month and 11 months aged groups in the neocortex (NEO), allocortex (ALLO), cerebellum (CB), olfactory bulbs (OB), medial basal forebrain (MF), hypothalamus (HYPO), brainstem (BS), cingulate cortex (CING), striatum (STR), ventral (VHIP) and dorsal hippocampus (DHIP). Data were analyzed using an adjusted F test with Kenward-Roger type adjustment of denominator degrees of freedom. n = 13 (NEO 12) for the group of 1-month-old rats and n = 13 (DHIP 12) for 11 months old rats. Statistical significance is indicated with *p ≤ 0.05, **p ≤ 0.01, ***p ≤ 0.001. The boxes span from the 25th to the 75th percentiles, the lines in the middle denote the median values, and the whiskers extend from the smallest values and up to the largest values (dots show all data points).

Since the changes were more significantly present in the a-series GSs, we decided to do an a to b-series ratio and identified the regions undergoing the most significant changes in the brain, which were - Ventral and dorsal hippocampus, medial forebrain, brainstem, and hypothalamus (Figure 5). There were no significant changes in the serum profile between the young and adult rats. However, in CSF, there was a slight decrease in GD1a.

**Figure 5.**
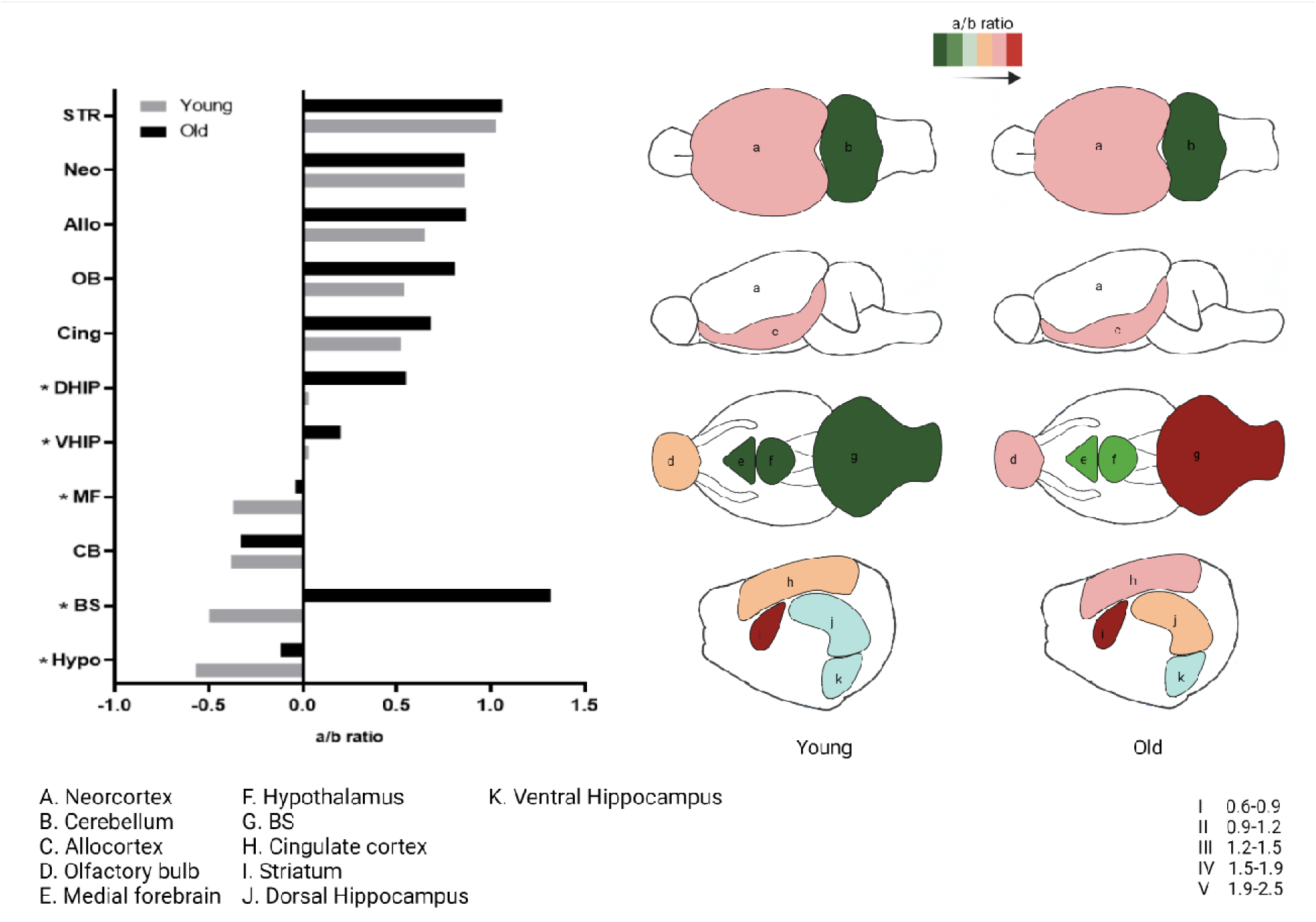
Bar chart depicting the a-series to b-series GSs in the brain regions. The a-series GSs included GM1, GM2, GD1a, and GM3, whereas the b-series included GD1b, GT1b, GQ1b, GD2, and GD3. Statistical significance is indicated with *p ≤ 0.05. The regions with the starkest changes during neurodevelopment were the hippocampus, hypothalamus, medial forebrain, and brainstem.

## Discussion

GSs play an essential role in functioning several brain cell types, most importantly, neurons. To study aging-related brain changes, a suitable model is imperative that could closely mimic human pathologies. Mouse models are frequently used as models of choice compared to rats in neuroscience^18^. However, rat models provide several physiological benefits along with genetic comparability with human traits^19^. They are bigger in size, and thus, convenient models requiring surgery or imaging. They are easier to handle, less stressed during contact with humans, and perform better in cognitive tests than mice^18^. Rats also possess similar number of tau isoforms as humans^20^. Hyperphosphorylation of tau is one of the hallmarks of AD, and thus, rat models could probably reflect tangle formation closely to humans than the mouse. Our study utilized the rat brain as a suitable translational model to study neurodevelopmental changes corresponding to aging.

The current study provided quantitative GS levels in several heterogeneous brain structures, along with the CSF and serum. Although lipidome of the rat brain has been studied previously, there is a lack of information on GSs and reports on changes in their levels through neurodevelopment. Our study was able to quantify nine major sub-class of GSs and observed and age and region-specific changes in GS profile.

In our study, we observed structures such as brain stem and cerebellum clustering together. Similar studies performed with proteome and metabolome in the rat brain have also shown clustering of these specific brain structures, demonstrating that functionally similar structures tend to have similar compositions of proteins and lipids ^10,12^. Interestingly, the rest of the structures were divided based on the presence of either predominantly b- or a-series GSs. The major brain GSs are distributed in specific locations and compositions in different cellular populations ^21^. GM1 is known to present in the myelin of the membrane, whereas the rest of the major GSs are known to present in the axolemma of the neurons^22^. GD1a, even though abundantly present of the neurons, is mainly present in the smaller neurons^23^. Our study observed regions such as striatum that are a copious amount of myelin to possess GM1 abundantly^24^. Another curious find was the distinct cluster of the ventral and dorsal hippocampus based on the GS levels. Although it is the same structure, there is functional heterogeneity within the different regions. The ventral hippocampus is more involved with spatial memory and learning, whereas the ventral hippocampus is more associated with emotional behaviour^25^. Thus, a detailed study can shed light on how a complex structure may have a different profile of GSs even though it is the same region morphologically.

The overall profile of GSs in the brain tissue alters with aging^26–29^. It is likely involved in neurodegeneration, leading to the development of diseases due to GSs composition’s influence on the neuronal membrane. Changes in the levels of lipids involved in GS syntheses such as ceramides and hexosylceramides have been previously reported. In general, most GS species’ content in the human brain decreases with age in normal subjects (age 25-85)^26^. Similar trends for the most abundant mono- (GM1a), di- (GD1a, GD1b), and trisialo (GT1b) species are observable in specific brain regions of AD patients (i.e., temporal and frontal cortex and nucleus basalis of Meynert) involved in the pathogenesis of the disease^30^. On the contrary, AD subjects showed an increased abundance of a-series species (i.e., GM2 and GM3) in the frontal and parietal cortex. Supposedly due to accelerated lysosomal degradation and astrogliosis occurring during the neuronal death^30^. The reduction of polysialoGSs (a-and b-series) and the increased monosialoGSs (a-series) content during aging was reported in the mouse and rat model of aging or accelerated senescence^31,32^. Specifically, GM1a was increased, and GD1a, GD1b, and GT1b were reduced. Another study demonstrated an increase of GM1a with age, significantly reduced absolute abundance of GD1a, GD1b, and GT1b^9^. Together with the increased enzymatic activity of catalytic neuraminidase (sialidase), the pattern was consistent with earlier reports^31,32^. Age-dependent reduction of glucocerebrosidase activity in the brain was reported in healthy individuals, especially substancia nigra and putamen^33^.

Neuraminidases are responsible for the enzymatic removal of sialic acids from sialoglycoconjugates (SGC’s). There are four different types in mammals - Neu1, Neu2, Neu3, and Neu4^34^. In neuroblastoma cells, they observed that Neu3 preferentially cleaved sialic acid present on the terminal position, which reduced the level of polysialated GSs, along with GM3 being converted to lactosylceramide. GSs such as GM1 and GM2 with a branching position sialic acid were resistant to degradation from the sialidase^35^. Neu3 activity in rat cerebellar granule cells increased with aging and increased β-galactosidase and β-Glucosidase activities and ceramide levels in the plasma membrane^36^.

There was also a change in a- to b-series ratio between young and adult rats, and some of the structures such as the brain stem and striatum were more affected than others. Thus, our study could potentially highlight the regions that might be initially susceptible to changes due to aging. However, our study could analyze only young and adult rats and the more significant changes of the aging are not visible here. Thus, more work is required to assess GSs levels in older rats and analyze other sub-classes of GSs.

## Methods

### Collection of cerebrospinal fluid

Rats were anesthetized with sodium pentobarbital intraperitoneally. The upper shoulder and the skull’s occipital areas were shaved and treated with an antiseptic solution (70% ethyl alcohol). A rat was placed onto a stereotactic frame, mounted with ear bars only, and put on a 3 cm height platform so that its front legs hung freely. The head was secured at an almost right angle with the rat’s body by gently pushing the stereotactic frame’s mouthpiece towards the nasal bone. The area between the occipital bone and the first cervical vertebra was located by gently touching the region through a sterile glove (approximately 2mm posterior from the interaural line in an adult rat), and an optimal spot for CSF withdrawal was determined. Using the stereotaxic micromanipulator, CSF was withdrawn with a collection apparatus consisting of a 23gauge needle connected with polyethylene tubing to a 1 ml syringe. New collection apparatus was used every time. The needle of the collection apparatus was aimed 5 mm ventrally from skin level. By gently pulling the syringe piston, slight negative pressure was created. If no CSF appeared in the tube, the needle was lowered by 0.5 mm, and the process was repeated until reaching 10 mm below skin level. CSF’s blood contamination was determined by checking for pink color in the samples of withdrawn fluid, and samples showing signs of contamination were discarded. When approximately 100-150 µl of CSF was collected, or no more CSF was possible to withdraw, the polyethylene tubing was cut off close to the collection needle using sterile scissors while simultaneously pulling the syringe piston to its upmost position. CSF was immediately transferred to 1.5 ml Eppendorf Protein LoBind tubes and snap-frozen in liquid nitrogen and stored at -80°C until processed for analysis.

### Collection of blood serum

Following the CSF collection, in anesthetized rats, the chest cavity was opened, and 1 ml of blood was collected from the heart’s left ventricle and transferred to 2 ml microcentrifuge tubes. After 15-30 min at room temperature, blood was centrifuged, the supernatant was collected, transferred to a new tube, snap-frozen in liquid nitrogen, and stored at -80°C until processed for analysis.

### Dissection of the brain tissue

Lastly, the brains were rapidly dissected from the skull, dropped into cold PBS for 30 seconds to cool down, and then placed on a reversed glass Petri dish resting on ice. The brains were partitioned into a set of 11 structures using a microsurgical knife (MSP Stab Knife Straight REF 7503, Surgical Specialties) and tweezers (Dumont #5B, World Precision Instruments). Structures in all brains were dissected in the same order. First, both olfactory bulbs were cut off at the level of the rostral pole of the hemisphere. Next, the cerebellum was separated by cutting it off at its joining with the brainstem. Medial basal forebrain, composed of the medial septum and the vertical limb of the diagonal band of Broca, was dissected as a tetrahedron-shaped piece of tissue with the wide base starting caudally at the optic chiasm and the sides following the border of the olfactory tubercles on both sides, coming together at their junction rostrally.

We chose to dissect the basal forebrain due to its particular role as the source of cholinergic projection neurons which are lost in Alzheimer’s disease^37^. Subsequently, the hypothalamus was dissected as a cylindrical piece of tissue starting rostrally at the optic chiasm, encircling the exposed hypothalamic surface and closing caudally just behind the mammillary bodies. The removed piece of tissue had 2-3 mm in height. Afterward, a midline cut through the corpus callosum was made, and the right hemisphere, including cortical and striatal tissue, was separated. A piece of the striatum, consisting primarily of caudate-putamen, was sampled devoid of an internal or external capsule’s fragments from this separated hemisphere. The hippocampal formation, including the dentate gyrus, CA3-CA1 fields, and the subiculum, was cut off at the hemisphere’s posterior margin. We partitioned the hippocampal formation into dorsal 2/3 and ventral 1/3 since these compartments have divergent neurochemical and functional specialization^25,38^. The remaining cortex was parcellated into three major parts. The cingulate cortex, a flap of cortex consisting of anterior cingulate and retrosplenial cortices, was dissected as a long strip of tissue cut along the hemisphere’s dorsal margin. The Allocortex, composed of the part of the cerebral cortex located below the rhinal sulcus, including the piriform cortex, entorhinal cortex, pallial amygdala, and associated transitional areas, was cut off at the rhinal sulcus. The remaining neocortex was sampled together with the underlying white matter. Finally, the brainstem was dissected just rostrally to the pontine nuclei, with the medulla oblongata trimmed at the joining with the spinal cord. All samples were immediately put into centrifugation tubes, buried in dry ice, and stored at -80°C.

### Mass spectrometry chemicals and reagents

Chemical standards of asialo-GM1 (cat. #ALX-302-013-M001), GM3 (cat. #ALX-302-005-M001), GD1a (cat no. ALX-302-007-M005), GD3 (cat. #ALX-302-010-M001), GT1b (cat. #ALX-302-011-M005), and GQ1b (cat. #ALX-302-012-MC05) were purchased from Enzo Life Sciences (Farmingdale, USA). GM1 standard (cat. #345724) was from Calbiochem (USA), GM2 (cat. #1502), and GD2 (cat. #1527) from Matreya (Pleasant Gap, PA). Solvents for the preparation of the UHPLC mobile phase for the dilution of stock solutions and sample extraction, i.e., LC-MS grade acetonitrile (cat. #001207802BS), methanol (cat. #0013687802BS), and 2-propanol (cat. #0016267802BS) were purchased from Biosolve BV (Dieuze, France), ammonium fluoride (cat. #338869) from Sigma-Aldrich (St. Louis, MO), chloroform (cat. #34854) and ammonium acetate (cat. #14267) from Honeywell (USA). Isotopically labelled [^13^C_18_]GM1 and [^13^C_18_]GM3 internal standards we prepared in-house as previously reported^39^.

### Brain tissue processing and extraction of GSs

Samples of dissected rat brain tissue (i.e., allocortex, brainstem, cerebellum, cingulate, dorsal hippocampus, hypothalamus, medial forebrain, neocortex, olfactory bulbs, striatum, ventral hippocampus) were received in Eppendorf tube and stored at -80 °C before lyophilization (8 h; ScanVac CoolSafe freeze dryers, LaboGene, Allerød, Denmark). Freeze-dried samples were powdered using one glass bead per sample on BeadBlasterTM 24 (Benchmark Scientific Inc.) homogenizer (speed 7 ms^-1^, time 2 s). Each sample of powdered tissue (∼2 mg) was accurately weighed. GSs were extracted from the powdered brain tissue using a slightly modified Folch protocol. The extraction protocol was as follows: initially, 1000 ul of chloroform/methanol (2:1, *v:v*) was added to ∼2 mg of each freeze-dried brain tissue sample and sonicated for 1 min; next 200 ul of deionized water was added, and the sample was further vortexed for 1 h. The sample was centrifuged (12000 rpm, 2 min), and 400 ul of the upper aqueous layer was collected into a clean Eppendorf tube. An aliquot of the supernatant (10 ul) was mixed with internal standards (51 ul) of [^13^]GM1 and [^13^C_18_]GM3 in 10 % 2-propanol, forming an internal standard concentration in a sample of 1.5 uM and 0.15 uM, respectively. 2 ul of each sample was injected in both positive and negative detection modes.

The extraction recovery and repeatability of above-mentioned extraction protocol was tested using isotopically labelled standard ([^13^C_18_]GM1 and [^13^C_18_]GM3) which were added to the pooled samples of allocortex brain tissue (Supplementary table 1).

### Processing and extraction of cerebrospinal fluid (CSF)

Rat CSF samples (20 – 40 ul) were received in Eppendorf tubes, stored at -80 °C and lyophilized for 8 h. Freeze-dried samples were re-dissolved in 20 ul of a mixture of [^13^C_18_]GM1 and [^13^C_18_]GM3 in 50 % 2-propanol, forming an internal standard concentration in a sample of 100 nM and 10 nM, respectively. The samples were vortexed and centrifuged (12000 rpm, 2 min). 2 ul of supernatant was injected onto the column using positive ion detection mode and 4 ul using negative ion detection mode.

### Processing and extraction of serum

50 ul of rat serum samples were stored at -80 °C in Eppendorf tubes and lyophilized for 8 h. Freeze-dried samples were powdered using one glass bead per sample in BeadBlasterTM 24 homogenizer (speed 6 ms^-1^, time 10 s). After addition of 5 ul of internal standards (mix [^13^C_18_]GM1 and [^13^C_18_]GM3 in 50 % 2-propanol forming an internal standard concentration in sample of 0.8 uM and 0.5 uM, respectively), GSs were extracted from the serum sample with 80 % 2-propanol (200 ul, sonicated for 1 min, 10 min of horizontal vortexing, 2000 rpm). The sample was centrifuged (12000 rpm, 2 min) and 170 ul of supernatant was collected into a clean Eppendorf tube. The pellet was then subjected to the same extraction procedure again by addition of 200 ul of 80 % 2-propanol. The supernatants from the first and second extractions were combined and dried under vacuum (Savant SPD121 P SpeedVac, Thermo Fisher Scientific, Inc.Waltham, USA). Once dry, the extracts were re-suspended in 50 ul of 50 % 2-propanol, vortexed and centrifuged (12000 rpm, 2 min). 2 ul of supernatant was injected into column using both positive and negative detection modes. In addition, in negative ion mode detection, the two times concentrated samples were analysed.

### Ultra-high performance liquid chromatography and mass spectrometry

Samples were analysed using ultra-high performance liquid chromatography (UHPLC), and tandem mass spectrometry (MS/MS) using selected reaction monitoring (SRM) technique. The UHPLC system (1290 Infinity II) and triple quadrupole (QqQ) mass analyser (model 6495) was from Agilent Technologies (CA, USA). We have used two different reversed-phase separations (for positive and negative ion detection mode) each carried on a C18 CSH analytical column (1.7 μm, 50 x 2.1 mm, cat. #186005296, Waters corp. UK) thermostated at 40 °C, at a flow rate of 0.3 ml/min. For positive ion mode detection, the mobile phase consisted from buffer A (water with 0.5 mM ammonium fluoride) and buffer B (2-propanol/methanol, 1:1, *v:v*) eluted in the linear gradient program as follows: 0.00 min 30 % B, 2.00 min 70 % B, 9.00 min 95% B, 13.00 min 95% B, 13.3 min 5% B, 14.3 min 5% B, 14.5 min 30% B, and 17.1 30% B. For negative ion mode detection, the mobile phase consisted from buffer A (10 mM ammonium acetate with 0.5 mM ammonium fluoride, pH=5.2) and buffer B (2-propanol/acetonitrile, 1:1, *v:v*) eluted in the linear gradient program as follows: 0.00 min 10% B, 4.00 min 85% B, 6.20 min 95% B, 10.20 min 95% B, 10.4 min 10% B, 14.4 min 95% B, 16.2 min 95% B, 16.40 min 10% B and 19.1 10% B. In both ion mode, injection volume was 2 ul and the following setting of QqQ mass analyser in ESI mode was used: capillary voltage of 3.5 kV (positive ion mode) and 3.0 kV (negative ion mode), gas flow rate 11 L/min at 130°C, sheath gas pressure 25 psi at 400 °C, nozzle voltage 500 V (positive ion mode) and 1500 V (negative ion mode). SRM libraries for both, positive and negative ion detection modes, were generated using Optimizer software (Agilent Technologies) from standard solution of an individual GSs. Up to 3 SRM transitions were used for a sufficient identification of GSs and a single best-performing SRM transition was used for quantification (Supplementary table 2).

### Quantification of GSs concentrations in samples

The calibration curves were generated adding isotopically-labeled [^13^]GM1 and [^13^C_18_]GM3 to determine the linearity range, limit of detection (LOD), and limit of quantification (LOQ). Matrix-matched (pooled of all samples of all brain structures) calibration curve was prepared in case of brain samples, at 6–8 concentration levels in the range of 9 – 9091 nM for [^13^C_18_]GM1 and 9.1 – 1364 nM for [^13^C_18_]GM3. Calibration curve for serum samples was prepared from pooled serum of 4 Wistar rats (provided by Department of Physiology, Masaryk University) at 7–8 concentration levels in the range of 25 – 500 nM for [^13^C_18_]GM3. In the case of CSF samples, calibration curves were prepared in 50 % 2-propanol solution, at 7–8 concentration levels in the range of 5 – 200 nM for [^13^C_18_]GM1 and 2.5 – 100 nM for [^13^C_18_]GM3. Each point of the dilution series was injected in triplicate, except for the lowest two calibration points measured in sextuplicate to accurately determine the standard deviation of integrated peak area and calculate LOD and LOQ as reported previously (Supplementary table 3, 4). The overall reproducibility of the GS quantitation was on average 15% (Supplementary table 5).

The quantification of GM1 and GM3 was performed by adding isotopically-labeled [^13^C_18_]GM1 and [^13^C_18_]GM3 to all analyzed samples. The concentration of other tested GSs detected in samples was semi-quantitatively determined using internal standardization with isotopically-labeled internal standard (IS) [^13^C_18_]GM3 in both positive and negative ion modes. The calculation is based on the concentration of IS added to samples, measured response (integrated peak area) of IS and response (integrated peak area) of GSs, and determined response factor (RF). RFs were determined in a conventional manner, and the mixture of GSs standard solutions was analyzed in triplicate in each sample set using the previously described gradient. Finally, the result was multiplied with a dilution factor derived from the relevant sample processing protocol. GS quantitation achieved in different sample matrices is listed in the Supplementary table 6.

### SRM data analysis

Quantitative results (i.e., integrated peak areas) were produced using commercial MassHunter Quantitative Analysis software (Agilent Technologies, La Jolla, USA) and further processed in Microsoft Excel (Microsoft Office Professional Plus 2016).

### Experimental design and statistical data analysis

The design of the experiment was based on the randomization of Latin Square Design (LSD). Statistical experimental design such as LSD minimizes the error in determining the parameters’ effect, allowing simultaneous, systematic, and efficient variation of all parameters than the classical method. They are used to increase power in particular situations, further taking account of the two sources of random effects. Continuous data are presented as means (SDs) if not otherwise stated. The continuous outcome variables characterizing GM1, GM2, GM3, GD1a, GD1b, GD2, GD3, GT1b, GQ1b volume (µg of GSs per mg of brain tissue DW) were analyzed with the use of a linear mixed model approach. The dichotomous covariates were gender (male vs. female), strain (Wistar vs. Sprague Dawley), and age (1-month vs. 11-month-old rats). Main outcomes were observed in 11 different brain areas (neocortex, allocortex, cerebellum, olfactory bulbs, medial basal forebrain, hypothalamus, brainstem, cingulate cortex, striatum, ventral and dorsal hippocampus). The main group effect age was estimated and means with confidence intervals and adjusted for gender and strain of an animal. In all models, the class variable “animal ID” was included as a repeated effect to correct accounting for animal subjectivity. The statistical test of the main effect was an adjusted F test with Kenward-Roger type adjustment of denominator degrees of freedom. The statistical evaluation of serum and CSF results was performed using an unpaired two-tailed t-test. Criterions for data exclusion from the statistical analysis were as follows: peak area of internal standard was under 45 % of internal standard mean peak area, and the detected value was under the LOQ, and data did not pass the test normality. SAS version 9.4 (SAS Institute Inc, Cary, NC, USA) was used for statistical analysis. Significance was accepted at the P ≤ 0.05 level. GraphPad Prism 8 version 8.3.0 (San Diego, CA, USA) was used for graphical expression.

## Author information

