## Supplemental Figure for "Brain map of aging-induced alterations in membrane ganglioside pattern"

**^6^** Waters corporation, Prague

**^7^** Faculty of Medicine, Masaryk University, Brno

**Supplementary Figure 1**

**Changes in the rat brain gangliosides levels dependent on various factors. a–c** Age, **d–g** gender, and **h–n** strain changes of ganglioside levels expressed as µg of gangliosides per mg of brain tissue DW. The difference between sum of all tested gangliosides (**a:** n = 1278, 1277; **d:** n = 1188, 1367; **h:** n = 1376, 1179) and individual gangliosides (**b:** n = 142, 141 for GQ1b and 142, 142 for others; **e:** n = 132, 151 for GQ1b and 132, 152 for others; **i:** n = 152, 131 for GQ1b and 153, 131 for others) in the whole brain, and between all tested gangliosides in individual brain structures (**c:** n = 108, 116 for NEO, 117, 108 for DHIP and 117, 117 for others; **f:** n = 108, 116 for NEO, 108, 108 for DHIP and 108,126 for others; **j:** n = 116, 108 for NEO, 126, 99 for DHIP and 126, 108 for others). g Gender-dependent difference of GQ1b levels in the hypothalamus (HYPO), n = 12, 14. **k** Effect of rat strain on GM2 in olfactory bulbs (OB) n = 14, 12, **l** GD3 in the medial basal forebrain (MF) n = 14, 12, **m** GM1, GM3, GQ1b in the brainstem (BS), n = 14, 12, and **n** GM1, GM2 in the cingulate cortex (CING), n = 14, 12. Data are shown as mean (S.D.) and evaluated by unpaired two-tailed t-test (**a–c**, **d–f**, and **h–j**) or by an adjusted F test with Kenward-Roger type adjustment of denominator degrees of freedom (**g**, and **k–c**). Statistical significance is indicated with *p ≤ 0.05, **p ≤ 0.01, ***p ≤ 0.001. Neocortex (NEO), allocortex (ALLO), cerebellum (CB), olfactory bulbs (OB), medial basal forebrain (MF), hypothalamus (HYPO), brainstem (BS), cingulate cortex (CING), striatum (STR), ventral (VHIP) and dorsal hippocampus (DHIP).

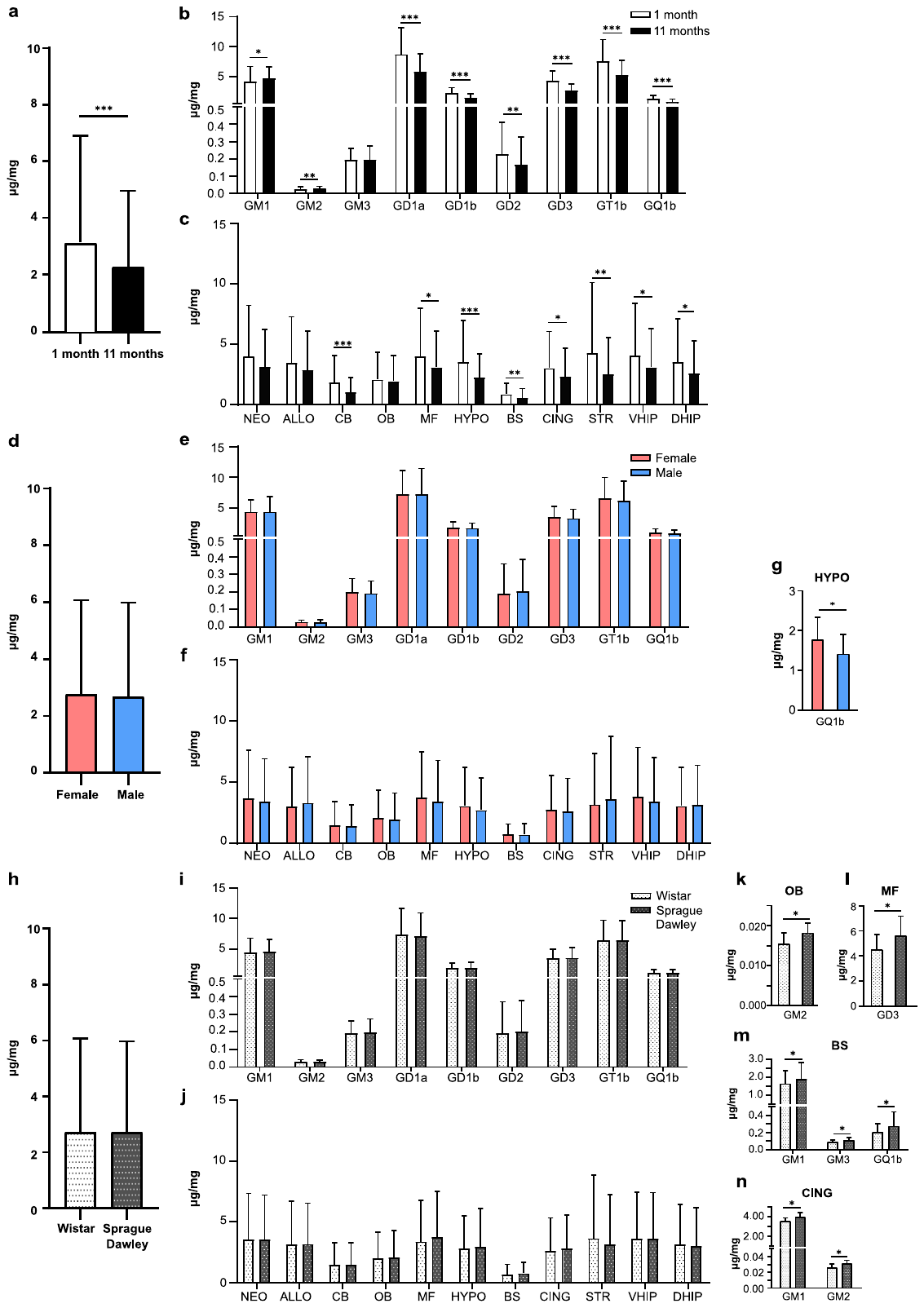
